# Genomic dissection of maternal, additive and non-additive genetic effects for growth and carcass traits in Nile tilapia

**DOI:** 10.1101/579334

**Authors:** Rajesh Joshi, John Woolliams, Theodorus Meuwissen, Hans Magnus Gjøen

## Abstract

**Background:** The availability of both pedigree and genomic sources of information for animal breeding and genetics has created new challenges in understanding how best they may be utilized and how they may be interpreted. This study computed the variance components obtained using genomic information and compared these to the variances obtained using pedigree in a population generated to estimate non-additive genetic variance. Further, the impact of assumptions concerning Hardy-Weinberg Equilibrium (HWE) on the component estimates was examined. The magnitude of inbreeding depression for important commercial traits in Nile tilapia was estimated for the first time, here using genomic data.

**Results:** The non-additive genetic variance in a Nile tilapia population was estimated from fullsib families and, where present, was found to be almost entirely additive by additive epistatic variance, although in pedigree studies this source is commonly assumed to arise from dominance. For body depth (BD) and body weight at harvest (BWH), the estimates of the additive by additive epistatic ratio (P<0.05) were found to be 0.15 and 0.17 in the current breeding population using genomic data. In addition, we found maternal variance (P<0.05) for BD, BWH, body length (BL) and fillet weight (FW), explaining approximately 10% of the observed phenotypic variance, which are comparable to the pedigree-based estimates. This study also disclosed detrimental effects of inbreeding in commercial traits of tilapia, which were estimated to cause 1.1%, 0.9%, 0.4% and 0.3% decrease in the trait value with 1% increase in the individual homozygosity for FW, BWH, BD and BL, respectively. The inbreeding depression and lack of dominance variance was consistent with an infinitesimal dominance model

**Conclusions:** An eventual utilisation of non-additive genetic effects in breeding schemes is not evident or straightforward from our findings, but inbreeding depression suggests for cross-breeding, although commercially this conclusion will depend on cost structures. However, the creation of maternal lines in Tilapia breeding schemes may be a possibility if this variation is found to be heritable.

## Background

This paper is a part of a wider study on the non-additive genetic effects in Nile tilapia and their potential utilization in tilapia breeding programs. A previous study [1] used the classical approach to partition the variance observed from a diallel mating design into additive, non-additive and maternal components using pedigree information to generate the additive and dominance relationship matrixes. These variance components are inferred from the variances within and between full-sib families, where the latter is also decomposed among sires and among dams.

These pedigree based selection methods have been gradually supplemented with, or replaced by, genomic information in various livestock species [2], and even in some commercial aquaculture species [3]. With the possibility of improved accuracy and more detailed information from genomics [4], there has been a growing interest to try to quantify and potentially utilize the non-additive genetic source of phenotypic variation. This new technology has introduced new challenges to fully understand the results of these methods and their equivalence to the classical decompositions based on pedigree. The availability of genomic information in Nile tilapia [5] has offered the opportunity to close this gap in an important aquacultural species. The first aim of this paper is to compare the genetic variance components obtained from using either genomics or pedigree information to generate the appropriate relationship matrices in a design generated to estimate non-additive variances.

The genomic BLUP (GBLUP) model builds a matrix of relationships between all individuals of a population based on genomic data, and BLUP uses these relations to partition the variance and predict the breeding values. The assumptions used to construct these relationship matrices have a direct effect on the accuracy of the results. There are different methods to construct the relationship matrices, most of them differing in the scaling parameters [6–8], which makes it difficult to make comparisons of results obtained with each of the methods. One method of comparison has been published by Legarra (2016) [9], where it is shown that re-scaling of the relationship matrices to the same reference population is necessary. In constructing relationship matrices, assumptions are often made about the presence of Hardy-Weinberg equilibrium (HWE), (e.g. in the use of Van Raden matrices [7], as used by GCTA [10]), and on managing the linkage disequilibrium (LD) [11]. These assumptions influence the orthogonality of the estimates of the variance components and hence the validity and generality of their biological interpretation. Thus, the second aim of this paper is to examine the impact of assumption of HWE on the relationship matrices and the consequences for the estimation.

Inbreeding depression is a natural phenomenon that is widely assumed to be deleterious for traits of commercial importance and thus has serious practical implications [12–15]. It has greater impact in populations with smaller effective population size (N_e_) than in those with higher N_e_, due to more efficient purging of deleterious alleles in the latter [16,17], which makes it a concern to breeders since N_e_ is often restricted in breeding populations. Genomic data allows a direct assessment of the extent of homozygosity and its variation rather than a reliance on changes predicted as a consequence of pedigree inbreeding. Consequently, utilisation of genomic data may contribute to a better design and operation of breeding programs. To date, the authors are unaware of estimates of inbreeding depression in Nile tilapia, even using the pedigree. Thus, the final aim of this paper is to quantify the effect of inbreeding depression for important commercial traits in Nile tilapia using genomic data.

Hence, this paper has tried to dissect the maternal, additive and non-additive genetic effects for growth and carcass traits in Nile tilapia, examining the impact of the assumption of HWE on the genomic relationship matrices and quantifying the inbreeding depression for these commercial traits.

## Methodology

### Experimental design, phenotypes and genotypes

The population used in this study and the experimental design have been previously described in more detail [1]. In short, the population was obtained from the reciprocal crossing of 2 parent groups, A and B, of Nile tilapia. The matings were partly factorial so that each parent used, male or female, had offspring that were both full-sibs and half-sibs. All offspring were hormonally treated, i.e. were either males or sex-reversed males, a normal aquacultural procedure to avoid sexual maturation, which may largely abrupt the growth, especially among females. Offspring were reared in three batches and harvested over 8 different days after 6-7 months in the grow-out ponds. The fish were filleted by three filleters. The phenotypes recorded were body weight at harvest (BWH), body depth (BD), body length (BL), body thickness (BT), fillet weight (FW) and Fillet yield (FY). Phenotypes were obtained on a total of 2524 individuals, with 1318 and 1206 from each of the two reciprocal crosses, in altogether 155 full-sib families.

From these, 1882 Nile tilapia samples were only genotyped using the Onil50 SNP-array (see Joshi et al. (2018) [5] for details). The raw dataset contained 58,466 SNPs, which were analysed using the Best Practices Workflow with default settings (sample Dish QC ≥ 0.82, QC call rate ≥ 97; SNP call-rate cutoff ≥ 97) in the Axiom Analysis Suite software [18]. Ten samples fell below the minimum QC call rate and were excluded. Then SNPs were selected based on the informativeness, i.e. based on the formation of clusters and resolution. Only SNPs classified as PolyHighResolution [18] (formation of three clusters with good resolution) and NoMinorHom [18] (formation of two clusters with no samples of one homozygous genotype) were selected, and 43,014 SNPs were retained. The mean SNP call rate for these SNPs was 99.5 (range: 97-100). Finally, SNPs were filtered for minor allele frequency (MAF ≥ 0.05), and 39,927 SNPs (68.3% of the total genotyped SNPs) were retained after all the quality control parameters had been applied. From the marker genotypes, the individual homozygosity was calculated as the proportion of homozygous loci per individual, and was incorporated into the models described below as a covariate for detecting directional dominance [19].

Of the 1882 genotyped, 1119 individuals from 74 full-sib families with an average of 15.1 offspring per full-sib family (range 1 to 44; standard deviation = 11.2) had phenotypic observation and were used for further analysis. Supplementary 1 shows the data structure and descriptive statistics in Tables S1.1 and S1.2 respectively, whereas scatterplots and the phenotypic correlations for these individuals are shown in Figure S1.1.

### Statistical Analysis

ASReml-4 [20] was used to fit mixed linear models, using REML to estimate variance components and breeding values. Eight different univariate GBLUP models were tested and compared for the six traits described above. The basic model used was an animal model (A), which was gradually expanded to an ADME (model with additive (A), dominance (D), maternal (M) and first order epistatic interactions (E) effects) by adding each effect as random effects in a heuristic approach. This resulted in the following models:

A model: **y = Xβ +h**b**+ Z_1_a + e**

AD model: **y = Xβ +h**b **+ Z_1_a + Z_2_d + e**

ADE model **y = Xβ +h**b **+ Z_1_a + Z_2_d + Z_3_e_aa_ + e**

ADME model **y** = **Xβ +h**b + **Z_1_a** + **Z_2_d** + **Z_3_e_aa_ + Z_4_m**+ **e**

ADM model: **y = Xβ +h**b **+ Z_1_a + Z_2_d + Z_4_m + e**

AM model: **y = Xβ +h**b **+ Z_1_a + Z_6_m + e**

AME model **y = Xβ +h**b **+ Z_1_a +Z_3_e_aa_ + Z_4_m + e**

AE model: **y = Xβ +h**b **+ Z_1_a + Z_3_e_aa_ + e**

where, **y** is the vector of records; **β** is the vector of fixed effects that account for reciprocal cross (1 d.f.), batch (2 d.f.) and day of harvest (7 d.f.); **h** the vector of overall marker homozygosity for each individual, with b the inbreeding depression parameter; **a** is a vector of random additive genetic effects; **d** is vector of random dominance effects; **e_aa_** is the vectors of first order additive x additive epistatic interactions; **m** is vector of maternal effects; **e** is a vector of random residual errors; and **X**, **Z_1_**, **Z_2_**, **Z_3_** and **Z_4_**, are corresponding design matrices for the fixed and random effects. For FW and FY, the fixed model also included filleter (2 d.f.). The (co)variance structures of the random effects are described below. Vectors **a**, **d**, **e_aa_** and **e** had effects for each individual having genotypes; **m** for each maternal family.

The models were also fitted with additive x dominance and dominance x dominance epistatic interaction effects, separately and in combination with additive x additive epistatic interaction effects. These parameters were bound to zero while solving the mixed model equations, thereby producing parameter values similar to those models without these random effects (results not shown). The distributional assumptions for the random effects were multivariate normal, with mean zero and

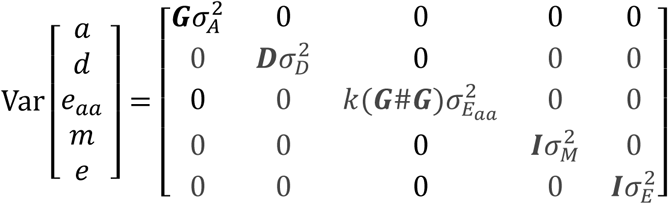

where 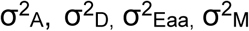 and 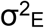 are additive genetic variance, dominance genetic variance, additive by additive epistatic variance, maternal variance and error variance respectively; **G** is the genomic relationship matrix with elements g_ij_; **D** is the dominance relationship matrix and **I** is an identity matrix of appropriate size. k(**G**#**G**) represents the additive by additive epistatic relationship matrix, where k is the scaling factor as described below and # is the Hadamard product of the two matrices given by 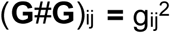 for elements in the indices i and j.

The phenotypic variance was calculated as 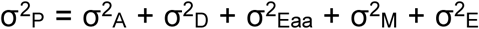, and the estimated variance components were expressed relative to the total phenotypic variance 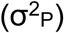: additive heritability 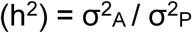, dominance ratio 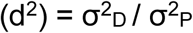 and maternal ratio 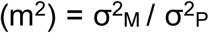. Broad sense heritability (H^2^) was calculated as 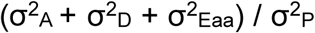 and the terms not in a model were set to 0. The variances obtained were also scaled by 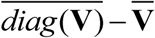 where **V** is their corresponding (co)variance matrix of size n and the bar denote the mean value [9].

Genomic natural and orthogonal interactions (NOIA) and Hardy-Weinberg Equilibrium (HWE) approaches were used to calculate the **G**, **D** and k(**G**#**G**) following the methods of [21]. These approaches differ in two ways: (i) the contrasts between genotypes used to define dominance deviations, and (ii) the scaling factors used for the relationship matrices.

The NOIA approach relaxes the assumption of HWE in the population, under which the genomic relationship matrix (**G**) is defined as:

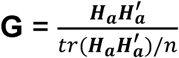

where, **H_a_** contains additive coefficients (*h*_*a*_) having the dimension of *n* × *m*, with *n* = number of animals and *m* = number of SNPs. *h*_*a*_ is coded as:

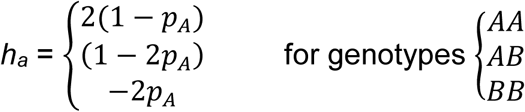

where, *p*_*A*_ is the frequency of allele *A*. For dominance deviations, NOIA uses the contrast that is orthogonal to h_a_ at each locus. Therefore, if p_AA_, p_AB_ and p_BB_ are the allelic frequencies of the respective genotypes, the dominance relationship matrix (**D**) is defined as;

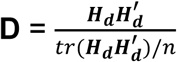

where, **H_d_** contains dominance coefficients (*h*_*d*_) defined for animal *i* and marker *j* by:

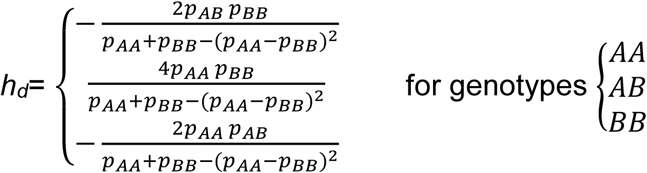

The epistatic relationship matrices were then calculated from the Hadamard projects and scaled using the average of the diagonals. Therefore, the additive by additive epistatic relationship was calculated as:

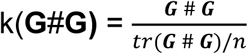

The HWE approach assumes that the population is under HWE equilibrium both in its scaling and in calculating the contrast for defining dominance deviations. If the locus is not in HWE the dominance contrast is not orthogonal to that for the additive effect, unlike in NOIA. The contrasts used to define the additive effects are unchanged but scaled assuming HWE, and the result is equivalent to method 1 of Raden [7]. So

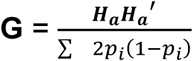

where the sum in the denominator is over all *m* loci. The dominance relationship matrix was calculated as

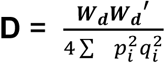

where **W_d_** contains elements *w*_*d*_ defined for animal *i* and marker *j*

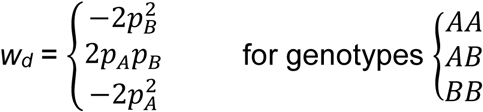

The scaling factor k for epistatic relationship matrices using the HWE approach was 1, so the additive by additive epistatic relationship matrix is simply the Hadamard product between the two matrices. The scatterplots for different relationship matrices are presented in Figure S1.3 and Figure S1.4 of Supplementary 1.

The software used to calculate the matrices [21] did not accept missing genotypes. As described above, 0.4% of genotypes were missing and these were predicted using R code [22] by sampling from {0,1,2} with the probabilities for each given by observed probabilities for that SNP. The effect of this prediction was checked with GCTA [10] by constructing the GRMs including and excluding the imputed genotypes. The correlation of >0.9995 between the additive and dominance relationships constructed using these two sets of genotypes suggest that there is no significant effect of prediction of the missing genotypes on our results as seen from the scatterplots of relationships in Figure S1.2 of Supplementary 1.

### Comparison of Models

Likelihood ratio tests were used to measure the goodness of fit for the models. The critical values were corrected for boundary effects following [23]. The critical values are obtained from a mixture of χ^2^ distributions with different degrees of freedom (d.f.) and were obtained for standard thresholds (P < 0.05, 0.01 and 0.001) by iteration using R. The distributions of the likelihood under the null hypothesis of zero variances for 1, 2 and 3 components were 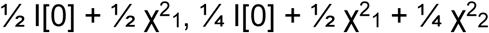 and 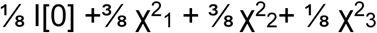 where I[0] corresponds to a point mass of 1 at x=0.

## Results

### Genetic architecture

The six traits could be differentiated into three distinct groups based on the scores of their likelihood ratio tests for the various models (Table 1): BD and BWH showed evidence of significant maternal environmental effects and non-additive genetic effects in the form of additive by additive epistasis. BL and FW showed evidence of significant maternal environmental effects only; whereas BT and FY showed no evidence of neither maternal environmental nor additive by additive epistatic effects. None of the traits showed significant dominance variance. The assumption of HWE in the breeding population did not influence the goodness of fit for any of the model, as the log likelihood values were identical. This is expected since the models are equivalent and only the parametrization differs.

**Table 1:** Log likelihood values with significance levels for different models for the six traits. The significance level for the likelihood ratio tests are expressed relative to the full model ADME. The critical values for Type 1 errors of 0.05, 0.01 and 0.001 were: for 1 d.f., 2.71, 5.42 and 9.55, respectively; for 2 d.f., 4.24, 7.29 and 11.77; and for 3 d.f. 5.44, 8.75 and13.48 respectively. The statistical significance is labelled as ‘*’, ‘**’ and ‘***’ for P<0.05, P<0.01 and P<0.001, respectively.

| Models | d.f. | BD | BL | BT | BWH | FW | FY |
| --- | --- | --- | --- | --- | --- | --- | --- |
| **ADME** |  | -43.48 | -191.28 | -1.78 | -31.51 | -69.90 | -68.55 |
| **ADE** | **1** | -46.55** | -195.75** | -2.25 | -35.82** | -74.74*** | -69.10 |
| **ADM** | **1** | -45.14* | -192.02 | -2.34 | -33.40* | -70.40 | -68.65 |
| **AME** | **1** | -43.48 | -191.28 | -1.78 | -31.51 | -69.90 | -68.55 |
| **AD** | **2** | -49.29** | -197.99*** | -3.04 | -39.29*** | -76.05*** | -69.25 |
| **AE** | **2** | -46.55* | -195.75** | -2.25 | -35.82** | -74.74** | -69.10 |
| **AM** | **2** | -45.15 | -192.02 | -2.40 | -33.40 | -70.40 | -68.65 |
| **A** | **3** | -49.29** | -197.99** | -3.06 | -39.29*** | -76.05** | -69.25 |

### Inbreeding depression

Detrimental effects of genomic homozygosity were evident for all of these commercial traits, although of different magnitudes. BWH and FW were found to be more sensitive to inbreeding than the other traits, with about 1% decrease in the trait value per 1% increase in the individual homozygosity (Table 2). The difference between upper and lower 5 percentile for homozygosity in this population was 0.062, and the resulting differences in performance were ∼6%, i.e. 23.21 g for BWH, 0.21 g for BD, 0.47 cm for BL and 9.76 g for FW. Traits BT and FY, the two traits with no evidence of non-additive genetic and maternal environmental effects, were found to be least sensitive with the estimates not differing significantly from 0 (P>0.05).

**Table 2:** Inbreeding depression for the commercial traits in Nile tilapia. “b” is the regression coefficient of trait on individual homozygosity, and D is the percentage decrease in the trait value per 1% increase in the individual homozygosity due to inbreeding depression. Standard errors are presented inside the parenthesis (). ** indicates p values 0.001 - 0.01 and * indicates p values 0.01 - 0.05 for significant values. “Difference” is the difference in performance between the upper and lower 5 percentile for homozygosity in the population. “Unit” is the unit for “b” and “Difference” of different traits.

|  | **BD** | **BWH** | **BL** | **FW** | **BT** | **FY** |
| --- | --- | --- | --- | --- | --- | --- |
| **b** | -3.27**<br>(1.19) | -371**<br>(137) | -7.57*<br>(2.95) | -156**<br>(56) | -7.08<br>(5.05) | -6.90<br>(4.93) |
| **D** | 0.37 | 0.91 | 0.34 | 1.08 | 0.17 | 0.21 |
| **Difference** | 0.21 | 23.22 | 0.47 | 9.76 | 0.44 | 0.43 |
| **Unit** | cm | g | cm | g | mm | % |

### Decomposition of variance components

Estimates of the variance components with the HWE and NOIA approaches for all the models and traits are presented graphically in Figure 1. The summary table for the models selected based on the likelihood ratio test are presented in Table 3.

**Table 3:** Components and their ratios with phenotypic variance for the models of best fit for different traits. Standard errors are presented in parentheses. The ratios are: narrow heritability h^2^, broad heritability H^2^, maternal ratio m2 and epistatic ratio 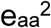

| Trait | Model | $\sigma^2_A$ | $\sigma^2_{Eaa}$ | $\sigma^2_m$ | $\sigma^2_e$ | $\sigma^2_p$ | $h^2$ | $H^2$ | $m^2$ | $e_{aa}^2$ |
| --- | --- | --- | --- | --- | --- | --- | --- | --- | --- | --- |
| **NOIA** |  |  |  |  |  |  |  |  |  |  |
| **BD** | AME | 0.086<br>(0.024) | 0.080<br>(0.049) | 0.047<br>(0.032) | 0.328<br>(0.044) | 0.541<br>(0.039) | 0.158<br>(0.042) | 0.307<br>(0.090) | 0.087<br>(0.055) | 0.148<br>(0.091) |
| **BWH** | AME | 699<br>(268) | 1183<br>(680) | 635<br>(418) | 4540<br>(618) | 7059<br>(498) | 0.099<br>(0.037) | 0.266<br>(0.093) | 0.090<br>(0.054) | 0.167<br>(0.096) |
| **BL** | AM | 0.284<br>(0.107) |  | 0.257<br>(0.162) | 2.803<br>(0.136) | 3.345<br>(0.209) | 0.085<br>(0.031) |  | 0.076<br>(0.045) |  |
| **FW** | AM | 118<br>(42) |  | 99<br>(63) | 1009<br>(50) | 1227<br>(79) | 0.096<br>(0.033) |  | 0.080<br>(0.047) |  |
| **BT** | A | 1.695<br>(0.441) |  |  | 8.015<br>(0.411) | 9.710<br>(0.458) | 0.174<br>(0.041) |  |  |  |
| **FY** | A | 1.758<br>(0.406) |  |  | 7.461<br>(0.378) | 9.220<br>(0.435) | 0.190<br>(0.039) |  |  |  |
| **HWE** |  |  |  |  |  |  |  |  |  |  |
| **BD** | AME | 0.097<br>(0.027) | 0.102<br>(0.063) | 0.047<br>(0.032) | 0.326<br>(0.045) | 0.573<br>(0.042) | 0.169<br>(0.046) | 0.348<br>(0.1) | 0.082<br>(0.053) | 0.178<br>(0.106) |
| **BWH** | AME | 791<br>(303) | 1504<br>(864) | 635<br>(418) | 4520<br>(626) | 7450<br>(544) | 0.106<br>(0.04) | 0.308<br>(0.104) | 0.085<br>(0.051) | 0.201<br>(0.111) |
| **BL** | AM | 0.321<br>(0.120) |  | 0.257<br>(0.162) | 2.801<br>(0.136) | 3.380<br>(0.213) | 0.095<br>(0.034) |  | 0.076<br>(0.044) |  |
| **FW** | AM | 133<br>(47) |  | 99<br>(63) | 1009<br>(50) | 1241<br>(81) | 0.107<br>(0.036) |  | 0.079<br>(0.047) |  |
| **BT** | A | 1.915<br>(0.498) |  |  | 8.004<br>(0.413) | 9.92<br>(0.492) | 0.193<br>(0.044) |  |  |  |
| **FY** | A | 1.987<br>(0.459) |  |  | 7.450<br>(0.379) | 9.437<br>(0.467) | 0.210<br>(0.042) |  |  |  |

**Figure 1:**
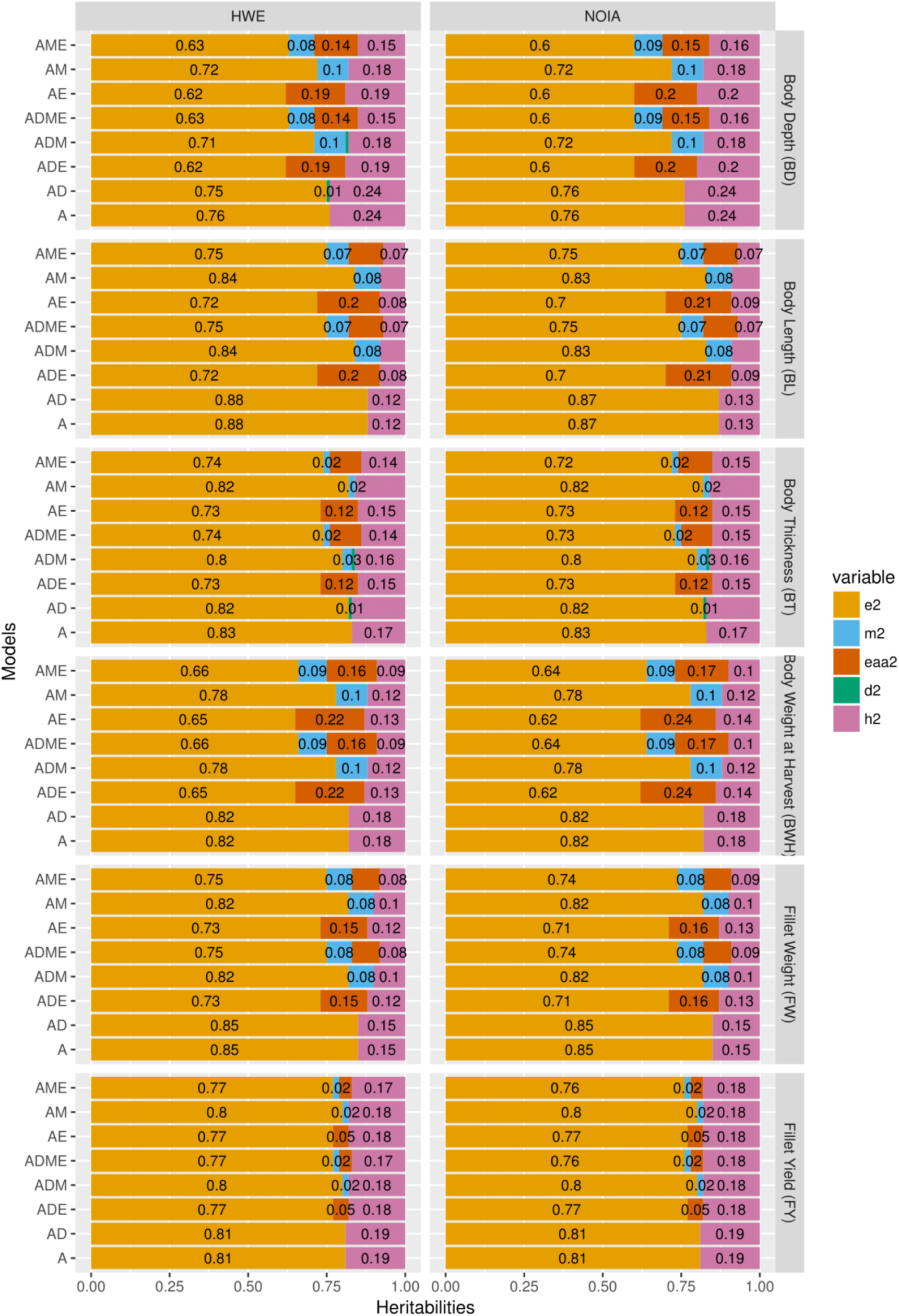
Decomposition of the phenotypic variance into different components using NOIA and HWE assumption approaches for the six traits. The ratios are: *h*^*2*^ is additive; *d*^*2*^ is dominance; 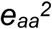 is additive by additive epistatic; *m*^*2*^ is maternal; and *e*^*2*^ is residual.

The simple A model gave the higher additive genetic variances, and the higher heritabilities across all the traits. Addition of dominance in the models had no effect on the estimated additive genetic variances, whereas including the additive by additive epistatic effect reduced the additive genetic variances markedly, except for BT and FY where there was no evidence (P>0.05) of epistasis. Inclusion of maternal environmental effects reduced the additive genetic variance compared to what was estimated with the simple A model, implying that without the maternal effect the additional variance associated with dams was interpreted as evidence of additive genetic effects. Including a maternal effect (AME models) also reduced the additive by additive epistatic variance compared to AE models. These reductions were again minimal for BT and FY. Similar results were obtained in both the NOIA and HWE assumption approaches. Hence, the numerical values are shown for the NOIA approach (scaled to the reference population [9]), unless otherwise mentioned.

Model dependent variation in the estimation of additive variance was also observed in the heritability estimates. For BT and FY, the two traits where the model of best fit was the simple A model, the heritabilities were least dependent on the models. For other traits, the differences observed among the models was up to 50%. For the best fit models, the estimates of the heritabilities were low to moderate, ranging from 0.08 ± 0.03 for BL to 0.19 ± 0.04 for FY (Table 4).

**Table 4:** Corrected heritabilities, ratio and variances for the models of best fit for different traits and approaches. The variances and ratios were corrected by (Mean (leading diagonal) – Mean) of the the corresponding relationship matrices as per Legarra (2016). Standard errors are presented in parenthesis.

| Traits | HWE |  |  |  | NOIA |  |  |  |
| --- | --- | --- | --- | --- | --- | --- | --- | --- |
| | $\sigma^2_A$ | $\sigma^2_{Eaa}$ | $h^2$ | $e_{aa}^2$ | $\sigma^2_A$ | $\sigma^2_{Eaa}$ | $h^2$ | $e_{aa}^2$ |
| **BD** | 0.086<br>(0.024) | 0.080<br>(0.049) | 0.159<br>(0.043) | 0.147<br>(0.091) | 0.086<br>(0.024) | 0.080<br>(0.049) | 0.159<br>(0.043) | 0.147<br>(0.091) |
| **BWH** | 698.774<br>(267.730) | 1169.547<br>(672.154) | 0.099<br>(0.037) | 0.167<br>(0.096) | 698.772<br>(267.729) | 1169.539<br>(672.149) | 0.099<br>(0.037) | 0.167<br>(0.095) |
| **BL** | 0.285<br>(0.107) |  | 0.085<br>(0.031) |  | 0.284<br>(0.107) |  | 0.085<br>(0.031) |  |
| **FW** | 117.948<br>(41.825548) |  | 0.096<br>(0.0324407) |  | 117.948<br>(41.825) |  | 0.096<br>(0.033) |  |
| **BT** | 1.694<br>(0.441) |  | 0.174<br>(0.041) |  | 1.694<br>(0.441) |  | 0.174<br>(0.041) |  |
| **FY** | 1.757<br>(0.406) |  | 0.191<br>(0.039) |  | 1.758<br>(0.406) |  | 0.191<br>(0.039) |  |

For BD and BWH, the traits for which the best fit model included additive by additive epistatic effect, the additive by additive epistatic ratio 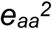 was 0.15 ± 0.09 and 0.17 ± 0.10 (Table 4), and additive by additive epistasis was found to be 48% and 63% of the total genetic variance for BD and BWH, respectively, but with large standard errors. Various other papers with genomic epistatic models also report large epistatic components [21,24,25] with corresponding large standard errors. Large differences between the individuals (Figure 2a) and the full-sib families (Figure 2b) were observed for the additive by additive epistatic effects.

**Figure 2:**
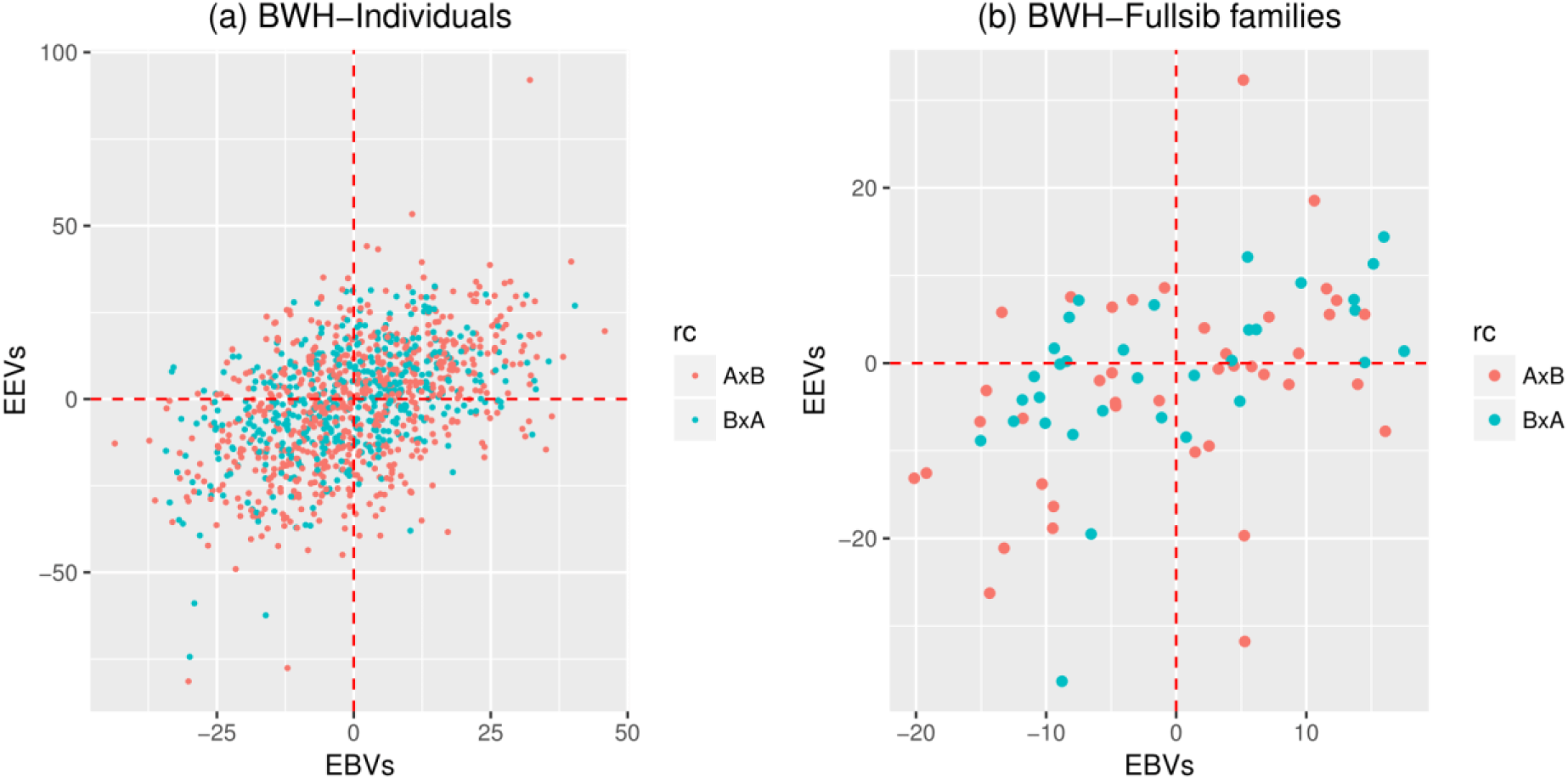
Scatterplot of estimated breeding values (EBVs) and epistatic (additive by additive) values (EEVs) for the trait BWH using NOIA approach (a) shows the scatterplot for all the individuals (b) shows the scatterplot for the mean values for different full-sib families. Please note that the values for x-axis and y-axis are different for both plots. The color of the dots in the scatterplot represents the types of reciprocal cross (rc): AxB and BxA.

For the four traits where the model of best fit included maternal environmental effect, the maternal ratio was found to be around 0.08±0.04 to 0.09± 0.06. As expected, this variance ratio was not affected by the two approaches or the models used. Thus, the previous recommendation [1] of possibility of creation of specialised maternal and sire lines in Nile tilapia breeding program is still relevant, if the maternal variance is found to be heritable.

## Discussion

### Interpretation of variance within the full-sib family

A major finding of this study is that the use of genomic relationship matrixes identified the source of non-additive genetic variance as being almost entirely additive by additive epistatic variance. The primary source of non-additive variance is commonly assumed to be dominance in pedigree based analyses [1,26,27], but this assumption can be very misleading as here, where the estimates of dominance variance were negligible. In this study, the information for estimating non-additive variance comes from the variance within full-sib families (see Supplementary Information 2), and in the presence of dominance and epistasis, the additional variance in full-sib families, above the additive variance provided by the sire and dam, is 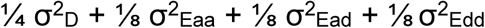 where 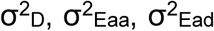 and 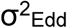 are dominance, additive by additive, additive by dominance and dominance by dominance epistatic variances [28]. Under an infinitesimal model with both additive and dominance effects, with the increase in the number of loci, either the dominance variance tends towards zero or the inbreeding depression tends towards infinity [28,29]. Thus, dominance may be present, but the genomic approach is showing this component behaves infinitesimally, with 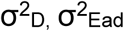 and 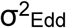 undetectable in analyses.

### Comparison with pedigree approach

This study adds a new dimension to our previous paper [1]. The availability of the genomic data in populations will inevitably lead to comparisons of genomic- and pedigree-based heritabilities, but these are not straightforward. Some publications argue that pedigree-based methods overestimate heritabilities [30–32], while some suggest the reverse [33–36], and other that the heritabilities are similar [37].

However, few studies recognize that the variance parameters obtained (i.e. the scaling parameters to the numerator or genomic relationship matrix) even in basic additive models do not refer to the same populations, and therefore the simple comparison of parameters can be rendered meaningless. For pedigree-based analyses the parameter refers to the base population of the pedigree (a subset of **A**), and for genomic-based analyses it can be viewed as the genetic variance in the population defined by the whole **G** assuming all the markers are in HWE. Many papers compare these values but they are uninformative as a large part of the difference can be accounted for by such distinctions [9,21]. To overcome the problem of comparability, the variance parameters from NOIA and HWE approaches were used to estimate the genetic variance in the entire population of this study [9] with marker genotypes as observed, equivalent to scaling the variance component estimates by 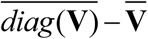, where **V** is the relevant relationship matrix and the bar denotes averaging elements.

In this study, where the models go beyond the additive components, there are additional reasons why components may differ. In the tilapia population studied here, the additive variance, when dominance is assumed to be the source of non-additive variation, gives a qualitatively different estimate to that obtained if additive epistasis is assumed (see Supplementary 2). Therefore, differences should be expected between the current study and [1]. A further issue with this study was that the data used was only a subset of the data used for [1], although Figure S1.5 of Supplementary 1 shows the sampling does not deviate far from random sampling expectations. This issue was overcome by repeating the pedigree analyses using only the phenotypes included in this study (see Table S1.4 in Supplementary 1).

The outcome from objective comparisons of the pedigree- and genomic analyses showed a qualitatively similar pattern of contributing sources of variance for all 6 traits insofar as additive, maternal and non-additive variances. Some small differences were observed: for example, the qualitative statistical significance for maternal ratio showed differences for BT and BL although the quantitative outcomes for the maternal ratio were similar. The evidence of non-additive genetic effects was found for the same traits (BD, BWH) irrespective of the type of relationships used. However, as mentioned above, critically, the genomics identified the source of non-additivity as additive by additive epistasis rather than dominance.

Genomic models were robust to misspecification in partitioning the variance among the components of the genetic and environmental models, and this robustness is another potential cause of difference between genomic and pedigree models. This is clearly observed when the basic model ‘A’ is fitted to traits for which the true genetic architecture is more complex (results are shown in Supplementary 1, Table S1.4). In the basic model ‘A’, using pedigree, the dam information is absorbed into estimating additive variance; in contrast to the genomic model, where it is the genotypes of the dam and its offspring that contribute information on the heritabilities, so the dam variance is no longer (wrongly) absorbed into the additive variance. Hence the pedigree-based heritabilities are higher for traits with maternal variance, as a consequence of the wrong model, and this difference was as large as 0.18.

### Impact of approaches used

GBLUP uses GRMs, and the assumptions in the construction of these GRMs can have a direct effect on the components; e.g. Van Raden matrices [7]) assume Hardy Weinberg equilibrium when scaling the relationship matrices, whereas this assumption is avoided with NOIA matrices. In this study, the use of these genomic approaches showed no difference to the qualitative outcomes related to the genetic architecture of the trait, but did make a quantitative difference e.g. additive-by-additive epistatic ratio 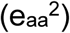 was inflated by ca. 20 % and 18%, and heritability (h^2^) by 6% and 10% for the traits BD and BWH respectively (Table 3). Such quantitative differences have also been observed in other studies [21]. As a consequence of the absence of dominance variance in this study, the differences between the NOIA and HWE collapse into differences in the scaling of the relationship matrices as the contrasts used to construct the matrices were identical. Therefore, the transformation of the components to a similar scale based on 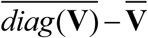 for these relationship matrices yielded identical variance components and ratios.

The NOIA and HWE approaches are statistical models in that they partition the variance observed in a population and use these parameters to estimate breeding values and dominance deviations [21]. As such, these estimates depend on the allele frequencies in the particular population, and the structure of the population which will influence the genotypic frequencies. A distinction needs to be made between the magnitudes of the variance components in the total genetic variance and the effects estimated using them on the one hand, and the ubiquity of the same phenomena in genotypic models (sometimes called biological models) on the other hand [38,39]. For example, the genotypes at a single locus may show complete dominance, but have a negligible dominance deviation, because the superior homozygote is very rare in the population. Although the NOIA approach removes limitations of HWE, there are major barriers to it moving towards the building of genotypic models. Firstly, it does not remove the impact of LD on estimates of the effects, and more seriously, the genotypic models are meaningful only if constructed with the causal variants and not with anonymous markers.

### Inbreeding depression

Absence of dominance variance does not necessarily mean the absence of inbreeding depression when the genetic architecture approaches the infinitesimal model, and evidence was found for depression in precisely the same four traits for which the basic ‘A’ model was rejected. To the authors’ knowledge, these estimates are the first for the commercial traits in Nile tilapia. Most of the quantification has been done using pedigree information in other aquaculture species, e.g. [40–42], and a few using genomics, e.g. [43]. In the present study, this information was not observable without the application of genomics because of the near identical inbreeding coefficients among individuals of the study population. Most of the traits clearly show the signal of inbreeding depression and ignoring this term leaves the estimates of the variance components and predictions of offspring merit open to bias (Supplementary 3). Further, the inbreeding depression is commercially significant for commercial traits, for example, FW decreases by 1% with 1% increase in homozygosity. Our population shows 6% difference between upper and lower 5 percentile for homozygosity in this population. This causes 6% difference for FW between individuals with high and low homozygosity, which has a huge commercial implication. Homozygosity can be minimized by controlling inbreeding, and by crossing unrelated lines. The latter will cause a large reduction in inbreeding depression if the regression on homozygosity holds also across lines.

In the infinitesimal model the allelic additive effects (*a’*) are if the order of 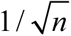 (i.e. *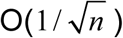*), as the number of loci, *n*, becomes large, so the additive variance remains finite. For inbreeding depression to remain finite the directional dominance deviations (d’) must be O(1/*n*), and so the consequence of an infinitesimal dominance model is that *d’*/*a’* must reduce by 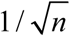 as *n* increases. This is consistent with biological pathway models such as [44], as when loci have increasingly small effect, responses will be more adequately described by the linear term based on the gradient of the response, and so the importance of partial dominance will diminish.

### Utilisation of the additive by additive epistatic effects

In the long run, additive by additive epistatic variance is expected to be exploited indirectly as it is converted to additive genetic variance due to random drift and selection; hence this form of variance affects the medium and long-term selection response indirectly [45]. Therefore, this argues for a simple breeding scheme, utilising only additive genetic effects, although re-structuring towards a cross breeding scheme, e.g. reciprocal recurrent selection, may be desirable for reasons related to the infinitesimal dominance detected or the inbreeding depression or the maternal variances.

Nevertheless, for some traits substantial additive by additive epistasis was observed even though it is expected that epistatic variance would be much smaller than the additive genetic variance in elite commercial populations [28,45]. This may prompt two questions. Firstly, whether these effects should be included in the estimation of genetic parameters: this is unlikely to be of benefit in selection decisions, partly because additive genetic variance already contains some of the variance arising from epistatic effects [24,28,46]. Secondly, whether the large epistatic ratio, predicting large differences among individuals in the population (Figure 2) can be used in the Nile tilapia breeding program in some way: since the observations of the epistasis relies upon anonymous loci, a more direct exploitation of epistasis will depend on finding out the causal variants showing large epistatic interactions [47,48] for different traits. This will require substantial resources to achieve, probably an order of magnitude greater than for identifying the additive effects of causal variants. Hence, this route seems rather complicated and costly to realise.

## Conclusion

This study has found that the non-additive genetic variance in the Nile tilapia population was almost entirely additive by additive epistatic variance, when using genomic relationship matrixes, whereas these non-additive effects are commonly assumed to be dominance using pedigree-based relationship matrixes. The inbreeding depression and lack of dominance variance was consistent with an infinitesimal dominance model. Finally, the creation of maternal lines in Tilapia breeding schemes may be a possibility if this variation is found to be heritable.

## List of abbreviations

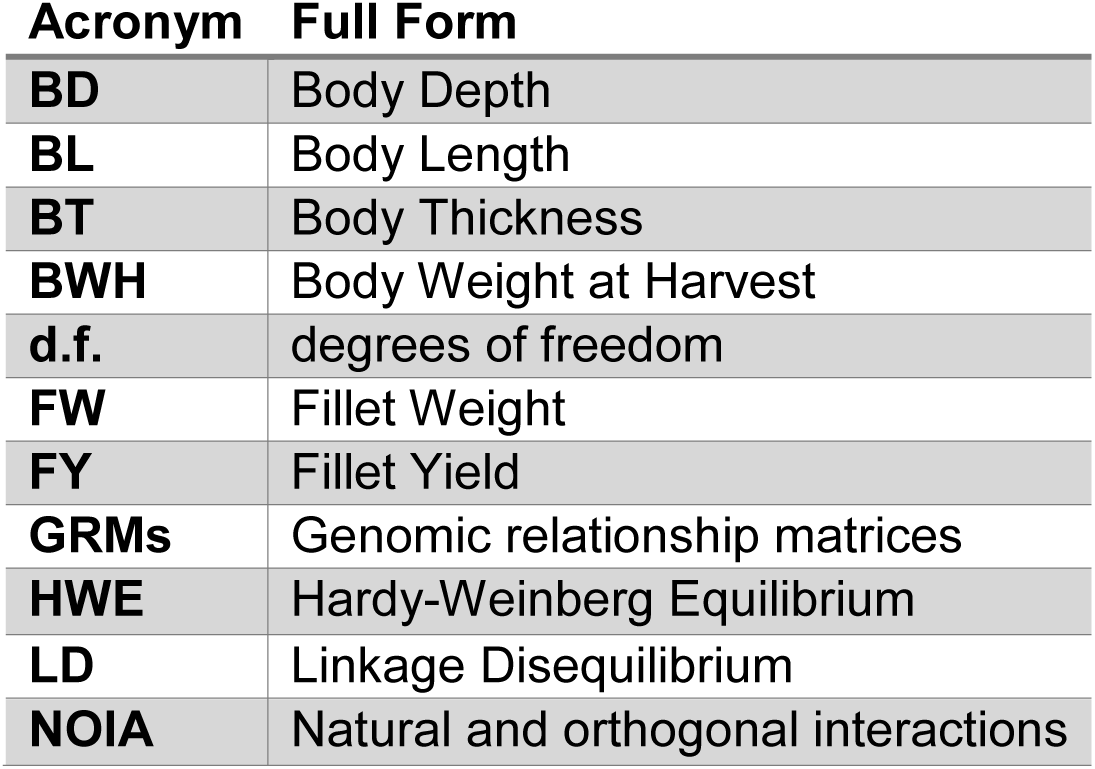

| Acronym | Full Form |
| --- | --- |
| **BD** | Body Depth |
| **BL** | Body Length |
| **BT** | Body Thickness |
| **BWH** | Body Weight at Harvest |
| **d.f.** | degrees of freedom |
| **FW** | Fillet Weight |
| **FY** | Fillet Yield |
| **GRMs** | Genomic relationship matrices |
| **HWE** | Hardy-Weinberg Equilibrium |
| **LD** | Linkage Disequilibrium |
| **NOIA** | Natural and orthogonal interactions |

## Declarations

### Ethics approval and consent to participate

Not applicable

### Consent for publication

Not applicable

## Availability of data and material

The genotype data used in the study are from commercial family material. This information may be made available to non-competitive interests under conditions specified in a Data Transfer Agreement. Requests to access these datasets should be directed to Alejandro Tola Alvarez.

## Authors’ contributions

HMG conceived and designed the study, RJ did the statistical analysis, JAW contributed to this analysis and developing the models used, and all authors contributed to the discussion of the results and writing of the paper.

### Acknowledgements

We would like to thank GenoMar AS for providing the data, in particular Anders Skaarud and Alejandro Tola Alvarez. JAW gratefully acknowledges funding by NMBU and by the UK Biotechnology and Biological Sciences Research Council BBS/E/D/30002275. ASReml analysis was performed in the Abel Cluster, Oslo.

## Authors’ information (optional)

## Supplementary files

Supplementary 1: Data Structure and the relationship matrices

Supplementary 2: Assumptions on the nature of non-additive genetic variance and the impact on estimates of additive genetic variance

Supplementary 3: Impact of inbreeding depression on models

## References

1. Joshi R, Woolliams J, Meuwissen T, Gjøen H. Maternal, dominance and additive genetic effects in Nile tilapia; influence on growth, fillet yield and body size traits. Heredity (Edinb) [Internet]. Nature Publishing Group; 2018 [cited 2018 Jan 16];1. Available from: http://www.nature.com/articles/s41437-017-0046-x

2. Meuwissen T, Hayes B, Goddard M. Genomic selection: A paradigm shift in animal breeding. Anim Front. American Society of Animal Science; 2016;6:6–14.

3. Hosoya S, Kikuchi K, Nagashima H, Onodera J, Sugimoto K, Satoh K, et al. Genomic Selection in Aquaculture. Bull Jap Fish Res Edu Agen No. 2017;45:35–9.

4. Gianola D, Fernando RL, Stella A. Genomic assisted prediction of genetic value with semi-parametric procedures. Genetics. Genetics Soc America; 2006;

5. Joshi R, Arnyasi M, Lien S, Gjoen HM, Alvarez AT, Kent M. Development and validation of 58K SNP-array and high-density linkage map in Nile tilapia (O. niloticus). bioRxiv. Cold Spring Harbor Laboratory; 2018;322826.

6. Speed D, Balding DJ. Relatedness in the post-genomic era: is it still useful? Nat Rev Genet. Nature Publishing Group; 2015;16:33.

7. VanRaden PM. Efficient Methods to Compute Genomic Predictions. J Dairy Sci. 2008;91:4414–23.

8. Yang J, Benyamin B, McEvoy BP, Gordon S, Henders AK, Nyholt DR, et al. Common SNPs explain a large proportion of the heritability for human height. Nat Genet [Internet]. 2010;42:565–9. Available from: http://dx.doi.org/10.1038/ng.608

9. Legarra A. Comparing estimates of genetic variance across different relationship models. Theor Popul Biol. Elsevier; 2016;107:26–30.

10. Yang J, Lee SH, Goddard ME, Visscher PM. GCTA: a tool for genome-wide complex trait analysis. Am J Hum Genet. Elsevier; 2011;88:76–82.

11. Speed D, Cai N, Johnson MR, Nejentsev S, Balding DJ, Consortium U. Reevaluation of SNP heritability in complex human traits. Nat Genet. Nature Publishing Group; 2017;49:986.

12. Fessehaye Y, Bovenhuis H, Rezk MA, Crooijmans R, van Arendonk JAM, Komen H. Effects of relatedness and inbreeding on reproductive success of Nile tilapia (Oreochromis niloticus). Aquaculture. Elsevier; 2009;294:180–6.

13. Christensen K, Jensen P, Jørgensen JN. A note on effect of inbreeding on production traits in pigs. Anim Sci. Cambridge University Press; 1994;58:298–300.

14. Smith LA, Cassell BG, Pearson RE. The effects of inbreeding on the lifetime performance of dairy cattle. J Dairy Sci. Elsevier; 1998;81:2729–37.

15. Bjelland DW, Weigel KA, Vukasinovic N, Nkrumah JD. Evaluation of inbreeding depression in Holstein cattle using whole-genome SNP markers and alternative measures of genomic inbreeding. J Dairy Sci. Elsevier; 2013;96:4697–706.

16. Meuwissen THE, Woolliams JA. Effective sizes of livestock populations to prevent a decline in fitness. Theor Appl Genet. Springer; 1994;89:1019–26.

17. Kristensen TN, Sørensen AC. Inbreeding–lessons from animal breeding, evolutionary biology and conservation genetics. Anim Sci. Cambridge University Press; 2005;80:121–33.

18. Thermo Fisher Scientific Inc. AxiomTMAnalysis Suite (AxAS) v4.0 USER GUIDE [Internet]. 2018. Available from: https://downloads.thermofisher.com/Affymetrix_Softwares/Axiom_Analysis_Suite_AxAS_v4.0_User_Guide.pdf

19. Xiang T, Christensen OF, Vitezica ZG, Legarra A. Genomic evaluation by including dominance effects and inbreeding depression for purebred and crossbred performance with an application in pigs. Genet Sel Evol [Internet]. BioMed Central; 2016 [cited 2017 May 15];48:92. Available from: http://www.ncbi.nlm.nih.gov/pubmed/27887565

20. Gilmour A, Thompson R. ASReml 4 Australasian Statistics Conference Port Lincoln 2014. 2014;

21. Vitezica ZG, Legarra A, Toro MA, Varona L. Orthogonal Estimates of Variances for Additive, Dominance, and Epistatic Effects in Populations. Genetics [Internet]. 2017 [cited 2018 Jan 29];206:1297–307. Available from: http://www.ncbi.nlm.nih.gov/pubmed/28522540

22. Mesfingo. R-code-for-missing-genotype-imputation [Internet]. 2016 [cited 2018 Jan 3]. Available from: https://github.com/Mesfingo/R-code-for-missing-genotype-imputation

23. Visscher PM. A note on the asymptotic distribution of likelihood ratio tests to test variance components. Twin Res Hum Genet [Internet]. Cambridge University Press; 2006;9:490–5. Available from: https://doi.org/10.1375/twin.9.4.490

24. Raidan FSS, Porto-Neto LR, Li Y, Lehnert SA, Vitezica ZG, Reverter A. Evaluation of non-additive effects in yearling weight of tropical beef cattle. J Anim Sci. 2018;

25. Piaskowski J, Hardner C, Cai L, Zhao Y, Iezzoni A, Peace C. Genomic heritability estimates in sweet cherry reveal non-additive genetic variance is relevant for industry-prioritized traits. BMC Genet. BioMed Central; 2018;19:23.

26. Shaw FH, Woolliams JA. Variance component analysis of skin and weight data for sheep subjected to rapid inbreeding. Genet Sel Evol. BioMed Central; 1999;31:1.

27. Hill WG. On estimation of genetic variance within families using genome-wide identity-by-descent sharing. Genet Sel Evol. BioMed Central; 2013;45:32.

28. Falconer DS, Mackay TF, Frankham R. Introduction to Quantitative Genetics (4th edn). Trends Genet. 1996;12:280.

29. Toro MA, Mäki-Tanila A. Some intriguing questions on Fisher’s ideas about dominance. J Anim Breed Genet. Wiley Online Library; 2018;135:149–50.

30. Haile-Mariam M, Nieuwhof GJ, Beard KT, Konstatinov K V, Hayes BJ. Comparison of heritabilities of dairy traits in Australian Holstein-Friesian cattle from genomic and pedigree data and implications for genomic evaluations. J Anim Breed Genet. Wiley Online Library; 2013;130:20–31.

31. Bérénos C, Ellis PA, Pilkington JG, Pemberton JM. Estimating quantitative genetic parameters in wild populations: a comparison of pedigree and genomic approaches. Mol Ecol. Wiley Online Library; 2014;23:3434–51.

32. Veerkamp RF, Mulder HA, Thompson R, Calus MPL. Genomic and pedigree-based genetic parameters for scarcely recorded traits when some animals are genotyped. J Dairy Sci [Internet]. 2011;94:4189–97. Available from: http://www.sciencedirect.com/science/article/pii/S002203021100422X

33. Tsai H-Y, Hamilton A, Tinch AE, Guy DR, Gharbi K, Stear MJ, et al. Genome wide association and genomic prediction for growth traits in juvenile farmed Atlantic salmon using a high density SNP array. BMC Genomics. BioMed Central; 2015;16:969.

34. Bangera R, Correa K, Lhorente JP, Figueroa R, Yáñez JM. Genomic predictions can accelerate selection for resistance against Piscirickettsia salmonis in Atlantic salmon (Salmo salar). BMC Genomics. BioMed Central; 2017;18:121.

35. Tan B, Grattapaglia D, Wu HX, Ingvarsson PK. Genomic relationships reveal significant dominance effects for growth in hybrid Eucalyptus. Plant Sci.Elsevier; 2018;267:84–93.

36. Zhang YD, Johnston DJ, Bolormaa S, Hawken RJ, Tier B. Genomic selection for female reproduction in Australian tropically adapted beef cattle. Anim Prod Sci. CSIRO; 2014;54:16–24.

37. Vitezica ZG, Varona L, Elsen J-M, Misztal I, Herring W, Legarra A. Genomic BLUP including additive and dominant variation in purebreds and F1 crossbreds, with an application in pigs. Genet Sel Evol [Internet]. BioMed Central; 2016 [cited 2016 Feb 2];48:6. Available from: http://gsejournal.biomedcentral.com/articles/10.1186/s12711-016-0185-1

38. Vitezica ZG, Varona L, Legarra A. On the additive and dominant variance and covariance of individuals within the genomic selection scope. Genetics [Internet]. 2013 [cited 2016 Jul 8];195:1223–30. Available from: http://www.ncbi.nlm.nih.gov/pubmed/24121775

39. Hill WG, Goddard ME, Visscher PM. Data and theory point to mainly additive genetic variance for complex traits. PLoS Genet. Public Library of Science; 2008;4:e1000008.

40. Rye M, Mao ILL. Nonadditive genetic effects and inbreeding depression for body weight in Atlantic salmon (Salmo salar L.). Livest Prod Sci [Internet]. Elsevier; 1998 [cited 2018 Sep 4];57:15–22. Available from: https://www.sciencedirect.com/science/article/pii/S0301622698001651

41. Pante MJR, Gjerde B, McMillan I. Effect of inbreeding on body weight at harvest in rainbow trout, Oncorhynchus mykiss. Aquaculture [Internet]. Elsevier; 2001 [cited 2018 Sep 4];192:201–11. Available from: https://www.sciencedirect.com/science/article/pii/S0044848600004671

42. Neira R, Díaz NF, Gall GAE, Gallardo JA, Lhorente JP, Manterola R. Genetic improvement in Coho salmon (Oncorhynchus kisutch). I: Selection response and inbreeding depression on harvest weight. Aquaculture [Internet]. Elsevier; 2006 [cited 2018 Sep 4];257:9–17. Available from: https://www.sciencedirect.com/science/article/pii/S0044848606001839

43. Hu G, Wang C, Da Y. Genomic heritability estimation for the early life-history transition related to propensity to migrate in wild rainbow and steelhead trout populations. Ecol Evol [Internet]. 2014 [cited 2015 Oct 16];4:1381–8. Available from: http://doi.wiley.com/10.1002/ece3.1038

44. Kacser H, Burns JA. Molecular democracy: who shares the controls? Biochem Soc Trans. Portland Press Limited; 1979;7:1149–60.

45. Hill WG. “Conversion” of epistatic into additive genetic variance in finite populations and possible impact on long-term selection response. J Anim Breed Genet. Wiley Online Library; 2017;134:196–201.

46. Mäki-Tanila A, Hill WG. Contribution of gene–gene interaction to genetic variation and its utilisation by selection. Proc 10th World Congr Genet Appl to Livest Prod. 2014.

47. Carlborg Ö, Jacobsson L, Åhgren P, Siegel P, Andersson L. Epistasis and the release of genetic variation during long-term selection. Nat Genet. Nature Publishing Group; 2006;38:418.

48. Große-Brinkhaus C, Jonas E, Buschbell H, Phatsara C, Tesfaye D, Jüngst H, et al. Epistatic QTL pairs associated with meat quality and carcass composition traits in a porcine Duroc× Pietrain population. Genet Sel Evol. BioMed Central; 2010;42:39.

