## Supplementary 1 for "Genomic dissection of maternal, additive and non-additive genetic effects for growth and carcass traits in Nile tilapia"

**Table S1.1:** Number of animals genotyped in different full-sib families. The row with P and G denotes the total number of phenotyped (coded black) and genotyped animals (coded bold blue) respectively from each full-sib family. Parents are coded from 1 to 18 in the header rows and columns. Empty cells means no phenotypes or genotypes were available for that full-sib family.

| | | **1** | **2** | **3** | **4** | **5** | **6** | **7** | **8** | **9** | **10** | **11** | **12** | **13** | **14** | **15** | **16** | **17** | **18** |
|---|---|---|---|---|---|---|---|---|---|---|---|---|---|---|---|---|---|---|---|
| **1** | **P** | | | | 5 | 9 | | 30 | | 5 | 5 | 26 | 26 | | 34 | | 15 | 35 | |
| | **G** | | | | | | | | | | | | | | | | | | |
| **2** | **P** | 30 | 20 | 18 | | | 11 | | 33 | | | | | 9 | | 3 | | | 16 |
| | **G** | **30** | **20** | **18** | | | **11** | | | | | | | **8** | | | | | |
| **3** | **P** | 29 | 18 | 25 | | | 6 | | 36 | | | | | 8 | | 13 | | | 36 |
| | **G** | | | | | | | | | | | | | | | | | | |
| **4** | **P** | 13 | 7 | 19 | | | 6 | | 23 | | | | | 11 | | 6 | | | 14 |
| | **G** | | | | | | | | | | | | | | | | | | |
| **5** | **P** | | | | | | | 6 | | 8 | 7 | 20 | 12 | | 25 | | 10 | 27 | |
| | **G** | | | | | | | **6** | | **8** | **7** | **20** | | | **21** | | **10** | **27** | |
| **6** | **P** | | | | 3 | 6 | | 46 | | 8 | 4 | 39 | 44 | | 59 | | 17 | 54 | |
| | **G** | | | | | | | **41** | | **8** | **4** | **39** | | | **38** | | **17** | **44** | |
| **7** | **P** | 32 | 9 | 17 | | | 8 | | 32 | | | | | 7 | | 2 | | | 30 |
| | **G** | **32** | **9** | **17** | | | **8** | | | | | | | **7** | | | | | |
| **8** | **P** | | | | 5 | 3 | | 24 | | 9 | 3 | 13 | 13 | | 19 | | 8 | 16 | |
| | **G** | | | | | | | **25** | | **8** | **3** | **13** | | | **10** | | **8** | **16** | |
| **9** | **P** | 29 | 16 | 26 | | | 9 | | 37 | | | | | 11 | | 3 | | | 27 |
| | **G** | **29** | **15** | **26** | | | **9** | | | | | | | **11** | | | | | |
| **10** | **P** | 35 | 27 | 16 | | | 8 | | 38 | | | | | 17 | | 6 | | | 30 |
| | **G** | | | | | | | | | | | | | | | | | | |
| **11** | **P** | | | | 1 | 2 | | 13 | | 2 | 1 | 8 | 13 | | 22 | | 8 | 21 | |
| | **G** | | | | | | | | | | | | | | | | | | |
| **12** | **P** | | | | 4 | 2 | | 36 | | 4 | 3 | 10 | 30 | | 47 | | 10 | 52 | |
| | **G** | | | | | | | **36** | | **4** | **3** | **10** | | | **34** | | **8** | **43** | |
| **13** | **P** | 19 | 3 | 14 | | | 3 | | 19 | | | | | 7 | | 1 | | | 10 |
| | **G** | **19** | **3** | **14** | | | **3** | | | | | | | **7** | | | | | |
| **14** | **P** | | | | 2 | 6 | | 14 | | | | 16 | 13 | | 15 | | 12 | 45 | |
| | **G** | | | | | | | **14** | | | | **16** | | | **9** | | **12** | **43** | |
| **15** | **P** | 16 | 15 | 22 | | | 6 | | 28 | | | | | 12 | | 4 | | | 29 |
| | **G** | **16** | **15** | **20** | | | **6** | | **1** | | | | | **12** | | | | | |
| **16** | **P** | 22 | 13 | 17 | | | 4 | | 26 | | | | | 15 | | 3 | | | 17 |
| | **G** | **22** | **13** | **17** | | | **5** | | **1** | | | | | **16** | | | | | **1** |
| **17** | **P** | | | | | 4 | | 17 | | 4 | 6 | 17 | 17 | | 34 | | 7 | 4 | |
| | **G** | | | | | | | | | | | | | | | | | | |
| **18** | **P** | | | | | | | 14 | | 6 | 3 | 12 | 22 | | 15 | | 8 | 18 | |
| | **G** | | | | | | | **14** | | **6** | **3** | **12** | **1** | | **13** | | **7** | **17** | |

**Table S1.2:** Descriptive statistics for the six traits, where N is the number of observation having both phenotypes and genotypes, SD is the standard deviation, SE is the standard error and CV is the coefficient of variation expressed as percentage.

| | N | Unit | Min | Max | Median | Mean (SE) | SD | CV% |
|---|---|---|---|---|---|---|---|---|
| **BWH** | 1119 | g | 115.60 | 802.80 | 390.20 | 407.31 (3.84) | 128.44 | 31.53 |
| **BL** | 1119 | cm | 14.10 | 28.00 | 22.40 | 22.38 (0.07) | 2.25 | 10.05 |
| **BD** | 1119 | cm | 5.00 | 12.00 | 8.80 | 8.89 (0.03) | 1.03 | 11.58 |
| **BT** | 1119 | mm | 12.90 | 59.70 | 40.50 | 40.70 (0.14) | 4.55 | 11.17 |
| **FW** | 1119 | g | 20.10 | 342.60 | 136.60 | 143.83 (1.56) | 52.32 | 36.38 |
| **FY** | 1119 | % | 15.24 | 50.53 | 33.15 | 32.83 (0.09) | 3.13 | 9.54 |

**Table S1.3:** Mean values of the genomic relationship matrices constructed with NOIA and HWE approaches

| | **HWE** | | | **NOIA** | | |
|---|---|---|---|---|---|---|
| | Overall | Diagonal | Off-diagonal | Overall | Diagonal | Off-diagonal |
| **G** | 0.00079 | 0.8847333 | -0.000791354 | 0.000893 | 1 | -0.000894454 |
| **D** | 0.038489 | 0.9250984 | 0.03690302 | 0.000893 | 1 | -0.000894454 |
| **k(G#G)** | 0.009216 | 0.7868253 | 0.007825131 | 0.011713 | 1 | 0.009945195 |
| **k(G#D)** | 0.002971 | 0.8211928 | 0.001507525 | 0.003311 | 1 | 0.001528032 |
| **k(D#D)** | 0.00513 | 0.8581108 | 0.003604039 | 0.004223 | 1 | 0.002441192 |

**Table S1.4:** Transformation of the variances on a similar scale based on the relationship matrices. The additive genetic variance ($\sigma^2_A$) and the heritability ($h^2$) were obtained from A model. No individual homozygosity was fitted in these models. The transformed variances and ratio are marked by * and was scaled by (Mean (diagonal) – Mean) of the the corresponding relationship matrices (Table S1.3) as per Legarra (2016)[1]. In "Ped", genomic relationship matrix was replaced by pedigree relationship matrix obtained using 3 generations of pedigree. The mean and mean(diagonal) of the relationship between 1119 individuals from pedigree relationship matrix were 0.2757587 and 1 respectively.

| Traits | Assumption | $\sigma^2_A$ | SE | $\sigma^2_A$* | SE | $h^2$ | SE | $h^{2*}$ | SE |
|---|---|---|---|---|---|---|---|---|---|
| BWH | NOIA | 1242 | 300 | 1241 | 300 | 0.18 | 0.04 | 0.19 | 0.04 |
| | HWE | 1404 | 339 | 1241 | 300 | 0.20 | 0.04 | 0.19 | 0.04 |
| | Ped | 3489 | 1391 | 2527 | 1007 | 0.44 | 0.14 | 0.36 | 0.10 |
| BD | NOIA | 0.13 | 0.03 | 0.13 | 0.03 | 0.24 | 0.04 | 0.24 | 0.04 |
| | HWE | 0.14 | 0.03 | 0.13 | 0.03 | 0.27 | 0.05 | 0.24 | 0.04 |
| | Ped | 0.31 | 0.12 | 0.23 | 0.09 | 0.50 | 0.15 | 0.42 | 0.11 |
| BL | NOIA | 0.40 | 0.12 | 0.40 | 0.12 | 0.13 | 0.04 | 0.13 | 0.04 |
| | HWE | 0.46 | 0.14 | 0.40 | 0.12 | 0.14 | 0.04 | 0.13 | 0.03 |
| | Ped | 1.20 | 0.50 | 0.87 | 0.36 | 0.33 | 0.12 | 0.26 | 0.08 |
| FW | NOIA | 177 | 47 | 177 | 47 | 0.15 | 0.04 | 0.16 | 0.04 |
| | HWE | 200 | 53 | 177 | 47 | 0.17 | 0.04 | 0.16 | 0.04 |
| | Ped | 536 | 215 | 388 | 156 | 0.39 | 0.13 | 0.32 | 0.09 |
| BT | NOIA | 1.70 | 0.44 | 1.69 | 0.44 | 0.17 | 0.04 | 0.18 | 0.04 |
| | HWE | 1.92 | 0.50 | 1.69 | 0.44 | 0.19 | 0.04 | 0.18 | 0.04 |
| | Ped | 1.83 | 0.92 | 1.32 | 0.67 | 0.18 | 0.08 | 0.14 | 0.06 |
| FY | NOIA | 1.76 | 0.41 | 1.76 | 0.41 | 0.19 | 0.04 | 0.19 | 0.04 |
| | HWE | 1.99 | 0.46 | 1.76 | 0.41 | 0.21 | 0.04 | 0.19 | 0.04 |
| | Ped | 2.65 | 1.17 | 1.92 | 0.85 | 0.26 | 0.10 | 0.21 | 0.07 |

[1] Legarra A. Comparing estimates of genetic variance across different relationship models. Theor Popul Biol. Elsevier; 2016;107:26–30.

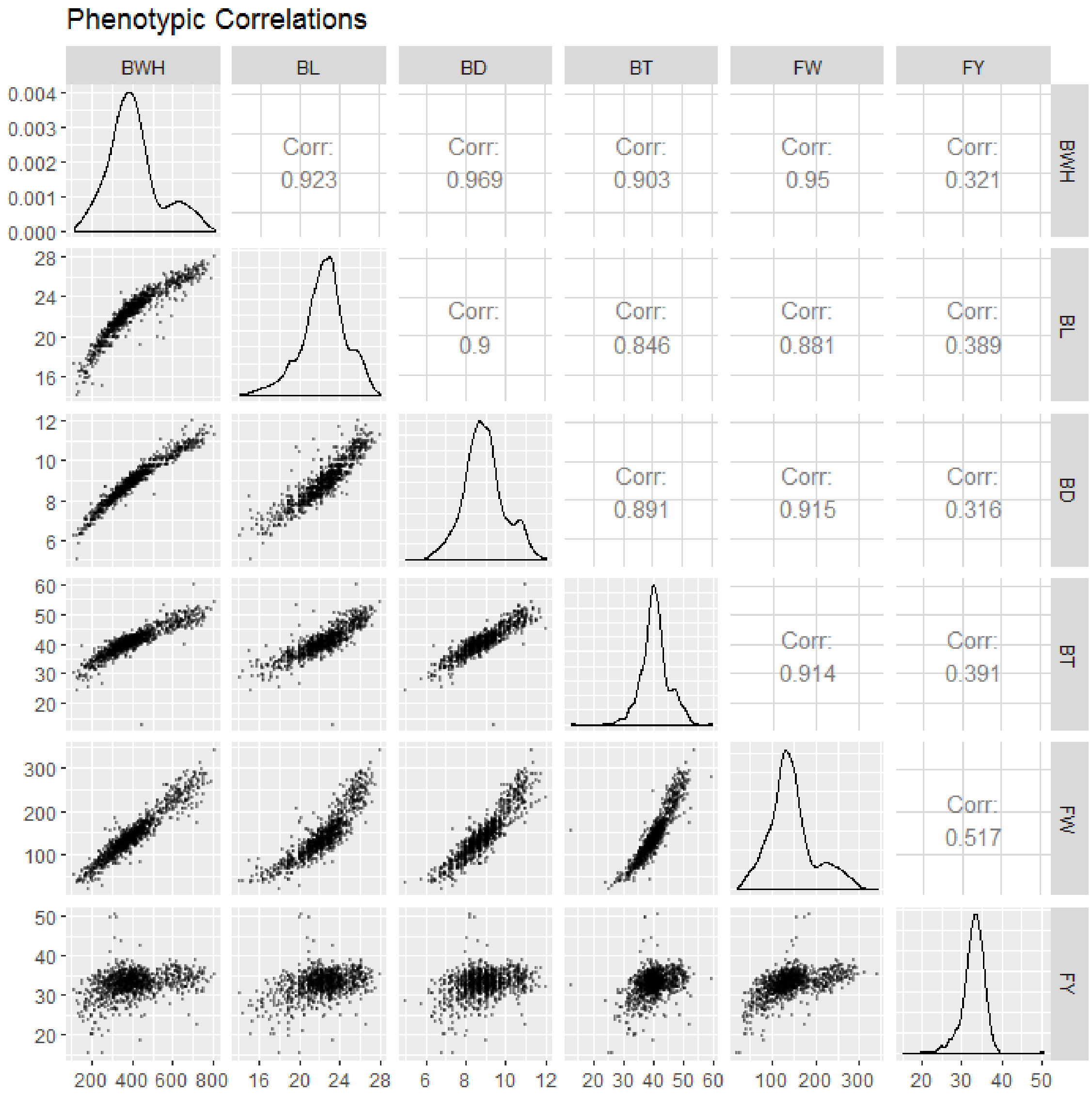


**Figure S1.1:** Scatterplots, histograms and the correlations of the 6 traits studied.

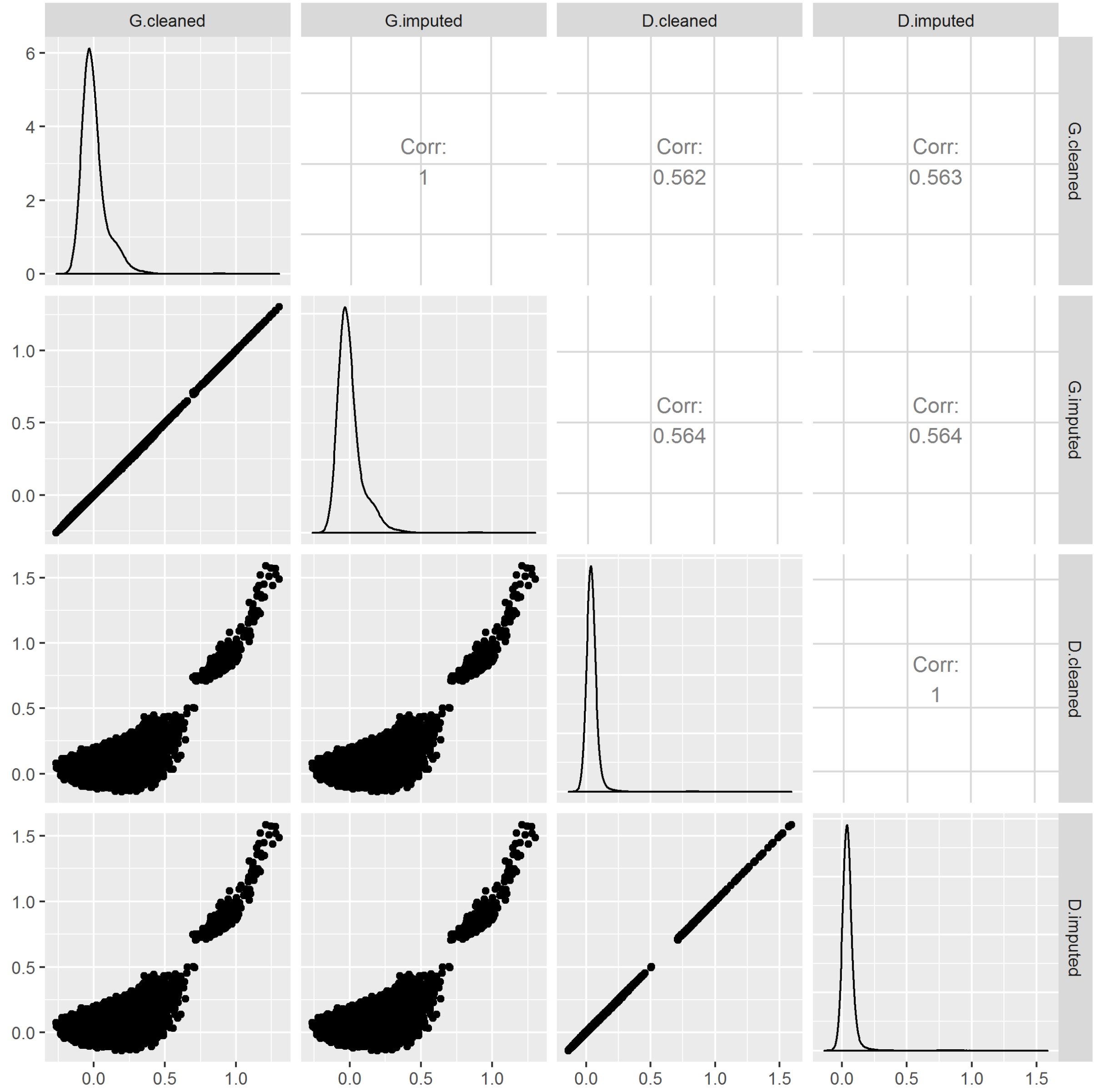


**Figure S1.2**: Scatterplot and correlation of the additive and dominance relationships using imputed genotypes (G.imputed and D.imputed respectively) and without imputed genotypes (i.e. with some missing genotypes, named as G.cleaned and D.cleaned respectively).

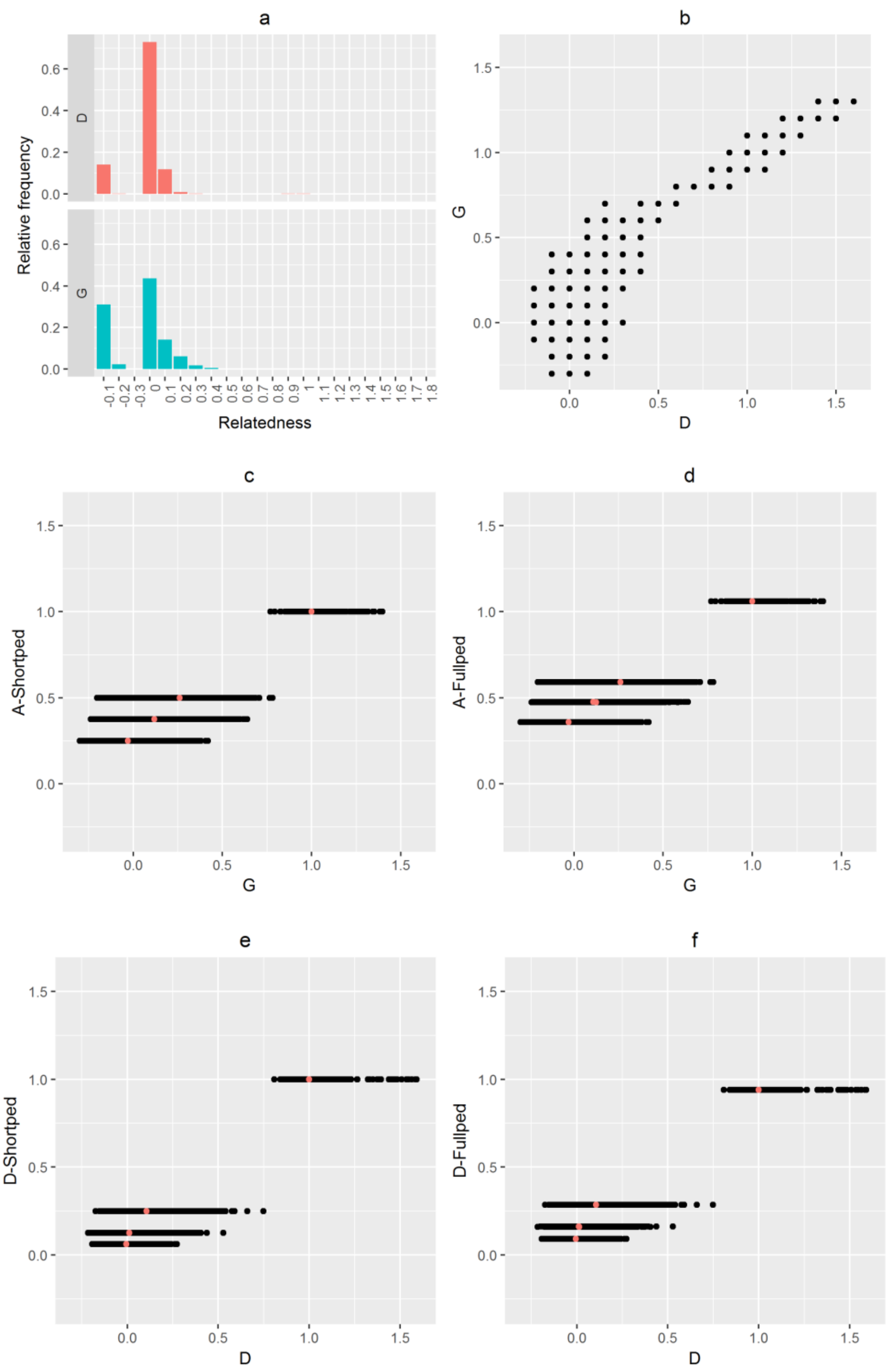


**Figure S1.3:** Scatterplots for different additive and dominance relationships. Mean of G and D are shown as orange dot in the respective plots **a)** Relative frequency plots for genomic relationships (G) and dominance relationships using genomic information (D). **b)** Scatterplot between G and D (correlation = 0.47). **c)** Scatterplot between G (using NOIA approach) and the pedigree relationships using 3 generations of pedigree (A-Shortped) (correlation = 0.72). **d)** Scatterplot between G and the pedigree relationships using 20 generations of pedigree (A-Fullped) (correlation = 0.72). **e)** Scatterplot between D and the dominance relationships using 3 generations of pedigree (D-Shortped) (correlation =0.65). **f)** Scatterplot between D and the dominance relationships using 20 generations of pedigree (D-Fullped) (correlation = 0.62).

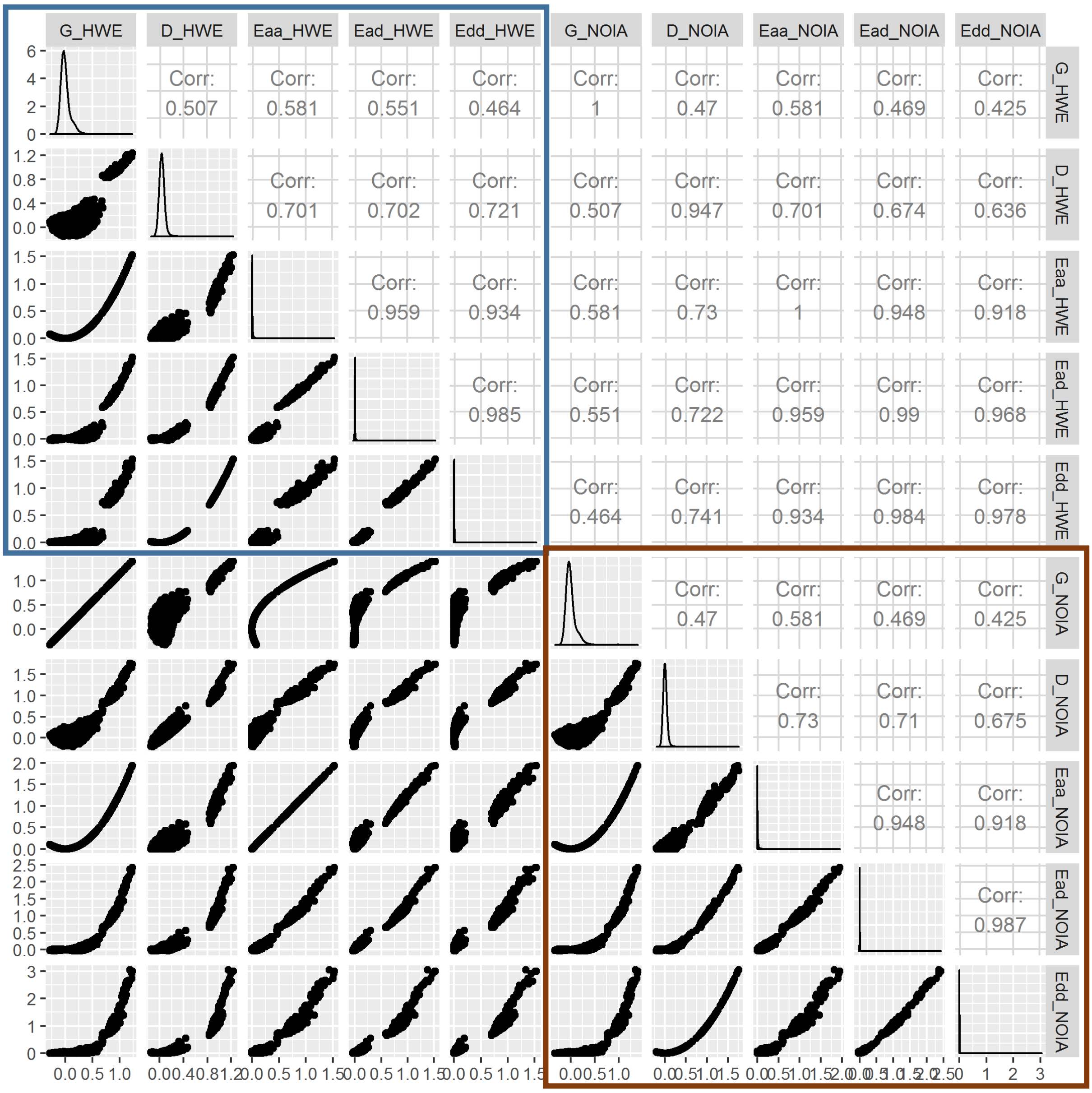


**Figure S1.4:** Scatterplots for different additive dominance and epistasis relationships using NOIA (inside brown box) and HWE assumption approaches (inside blue box).

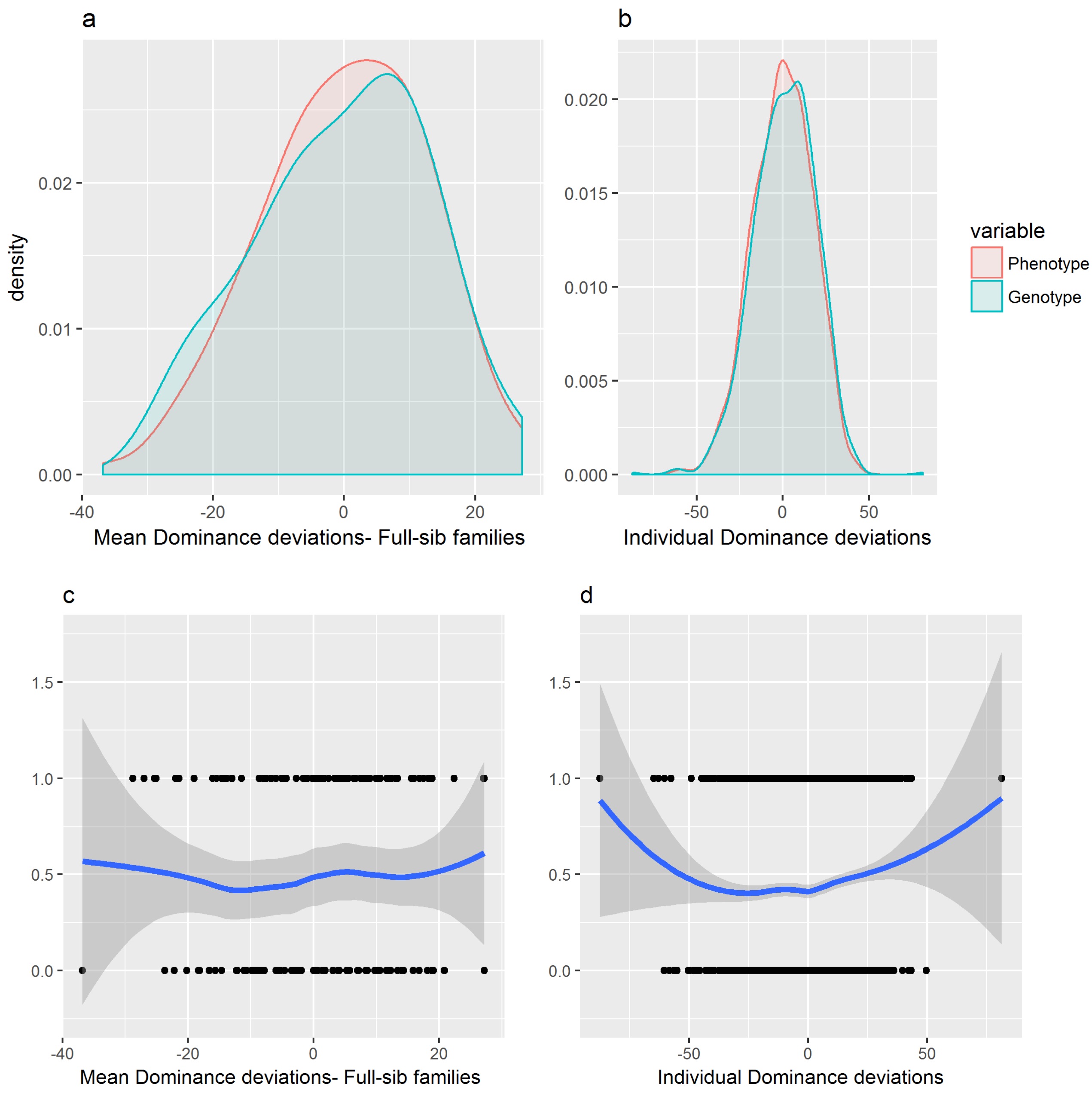


**Figure S1.5**: **a)** and **b)** Density plots showing the dominance deviations for BWH between the animals having phenotypes and those that were selected from them to be genotyped. **c)** and **d)** Scatterplot and LOESS regression between the selected and non-selected individuals. The selected individuals were coded as 1 and the non-selected individuals were coded as 0. The dominance deviations were obtained from the ADM model given in Joshi et al. (2018)[2].

---

[2] Joshi R, Woolliams J.A., Meuwissen T.H.E., Gjøen H.M.. Maternal, dominance and additive genetic effects in Nile tilapia; influence on growth, fillet yield and body size traits. *Heredity (Edinb)* [Internet]. Nature Publishing Group; 2018 Jan 16 [cited 2018 Jan 16];1. Available from: http://www.nature.com/articles/s41437-017-0046-x
