## Supplementary 2 for "Genomic dissection of maternal, additive and non-additive genetic effects for growth and carcass traits in Nile tilapia"

### Assumptions on the nature of non-additive genetic variance and the impact on estimates of additive genetic variance

Joshi et al. (2018)[1] showed how the variance components of the basic factorial mating design used in this study were translated into estimates of maternal, dominance and additive variances related to a pedigree. The basic factorial mating design had 3 core components ($V_{Sire}$, $V_{Dam}$, and $V_{Fsib}$) which can be related to the covariances (C) between individuals, *i* and *j*, assuming a mean is fitted to the population.

| | | | |
|---|---|---|---|
| *i*, *j* no common parent (U), | $C_U$ | $= 0$ | (1) |
| *i*, *j* paternal half-sibs (PHS), | $C_{PHS}$ | $= V_{Sire}$ | (2) |
| *i*, *j* maternal half-sibs (MHS), | $C_{MHS}$ | $= V_{Dam}$ | (3) |
| *i*, *j* full-sibs (FS), | $C_{FS}$ | $= V_{Sire} + V_{Dam} + V_{Fsib}$ | (4) |

For this population *i* and *j* were in generation 22, and Joshi et al. (2018)[1] published the main results with a base set at generation 20.

*Dominance*. Assuming the non-additive genetic variation was primarily arising from dominance then Joshi et al. (2018)[1] showed:

$$C_U = (4\sigma^2_A + \sigma^2_D)/16 \quad (5)$$

$$C_{PHS} = (6\sigma^2_A + 2\sigma^2_D)/16 \quad (6)$$

$$C_{MHS} = (6\sigma^2_A + 2\sigma^2_D)/16 + \sigma^2_M \quad (7)$$

$$C_{FS} = (8\sigma^2_A + 4\sigma^2_D)/16 + \sigma^2_M \quad (8)$$

The fitted mean will account for the genotypic drift from the base generation, which is represented by $C_U$, and Equation 5 can be subtracted from the (6), (7) and (8).

$$C_{PHS} = (2\sigma^2_A + \sigma^2_D)/16 \quad (9)$$

$$C_{MHS} = (2\sigma^2_A + \sigma^2_D)/16 + \sigma^2_M \quad (10)$$

$$C_{FS} = (4\sigma^2_A + 3\sigma^2_D)/16 + \sigma^2_M \quad (11)$$

[1] Joshi R, Woolliams J.A., Meuwissen T.H.E., Gjøen H.M.. Maternal, dominance and additive genetic effects in Nile tilapia; influence on growth, fillet yield and body size traits. *Heredity (Edinb)* [Internet]. Nature Publishing Group; 2018 Jan 16 [cited 2018 Jan 16];1. Available from: http://www.nature.com/articles/s41437-017-0046-x

Solving these equations and equating them to (2) to (4) results in the following:

- $\sigma^2_M$ is estimated as $C_{MHS} – C_{PHS}$ .
- $\sigma^2_D$ is estimated as $16(C_{FS}–C_{PHS}–C_{MHS}) = 16V_{Fsib}$
- $\sigma^2_A$ is estimated as $16C_{PHS}-8(C_{FS}-C_{MHS}) = 8(V_{Sire} - V_{Fsib})$.

*Epistasis*[2]. Consider now an assumption that the non-additive genetic variation was primarily arising from A#A, denoted $\sigma^2_I$ :

$$C_U = (16\sigma^2_A + 4\sigma^2_I)/64 \quad (5)$$

$$C_{PHS} = (24\sigma^2_A + 9\sigma^2_I)/64 \quad (6)$$

$$C_{MHS} = (24\sigma^2_A + 9\sigma^2_I)/64 + \sigma^2_M \quad (7)$$

$$C_{FS} = (32\sigma^2_A + 16\sigma^2_I)/64 + \sigma^2_M \quad (8)$$

As with dominance the fitted mean removes $C_U$ and this is subtracted from remaining covariances.

$$C_{PHS} = (8\sigma^2_A + 5\sigma^2_I)/64 \quad (6)$$

$$C_{MHS} = (8\sigma^2_A + 5\sigma^2_I)/64 + \sigma^2_M \quad (7)$$

$$C_{FS} = (16\sigma^2_A + 12\sigma^2_I)/64 + \sigma^2_M \quad (8)$$

The solutions to these equations are:

- $\sigma^2_M$ is estimated as $C_{MHS} – C_{PHS}$ .
- $\sigma^2_I$ is estimated as $32(C_{FS}–C_{PHS}–C_{MHS}) = 32V_{Fsib}$
- $\sigma^2_A$ is estimated as $28C_{PHS}-20(C_{FS}-C_{MHS}) = 8V_{Sire} - 20V_{Fsib} = 8(V_{Sire} - V_{Fsib}) – 12V_{Fsib}$.

Therefore the estimate of $\sigma^2_A$ from this design is reduced when the non-additive variation is assumed to be additive-by-additive epistasis rather than dominance, and this reduction is of the order of 3/8 $\sigma^2_I$.

[2] Cockerham CC. An extension of the concept of partitioning hereditary variance for analysis of covariances among relatives when epistasis is present. *Genetics.* Genetics Society of America; 1954;39(6):859.
