## Supplementary 3 for "Genomic dissection of maternal, additive and non-additive genetic effects for growth and carcass traits in Nile tilapia"

Both the models with HWE and NOIA approaches were fitted without individual homozygosity as the covariate to account for the impact of the inbreeding depression in the models. The summaries of the variance parameters are presented in the tables below.

**Table S3.1**: Heritabilities, ratio and phenotypic variance, for the models of best fit for different traits. The relationship matrices were constructed with HWE approach[1]. Models were not fitted with individual homozygosity as the covariate.

| HWE approach - Without individual homozygosity | | | | | | | | | | | |
|---|---|---|---|---|---|---|---|---|---|---|---|
| **Traits** | **Model** | **$h^2$** | **se** | **$e_{aa}^2$** | **se** | **$H^2$** | **se** | **$m^2$** | **se** | **phenvar** | **se** |
| **BD** | AME | 0.17 | 0.05 | 0.19 | 0.11 | 0.36 | 0.10 | 0.08 | 0.05 | 0.58 | 0.04 |
| **BWH** | AME | 0.11 | 0.04 | 0.22 | 0.11 | 0.33 | 0.10 | 0.08 | 0.05 | 7540 | 548 |
| **BL** | AM | 0.10 | 0.03 | | | | | 0.08 | 0.05 | 3.41 | 0.22 |
| **FW** | AM | 0.11 | 0.04 | | | | | 0.08 | 0.05 | 1252 | 82 |
| **BT** | A | 0.20 | 0.04 | | | | | | | 9.96 | 0.50 |
| **FY** | A | 0.21 | 0.04 | | | | | | | 9.45 | 0.47 |

**Table S3.2**: Heritabilities, ratio and phenotypic variance, for the models of best fit for different traits. The relationship matrices were constructed with NOIA approach[1]. Models were not fitted with individual homozygosity as the covariate.

| NOIA approach - Without individual homozygosity | | | | | | | | | | | |
|---|---|---|---|---|---|---|---|---|---|---|---|
| **Traits** | **Model** | **$h^2$** | **se** | **$e_{aa}^2$** | **se** | **$H^2$** | **se** | **$m^2$** | **se** | **phenvar** | **se** |
| **BD** | AME | 0.16 | 0.04 | 0.16 | 0.09 | 0.32 | 0.09 | 0.08 | 0.05 | 0.54 | 0.04 |
| **BWH** | AME | 0.10 | 0.04 | 0.19 | 0.10 | 0.29 | 0.09 | 0.09 | 0.05 | 7110 | 499 |
| **BL** | AM | 0.09 | 0.03 | | | | | 0.08 | 0.05 | 3.38 | 0.22 |
| **FW** | AM | 0.10 | 0.03 | | | | | 0.08 | 0.05 | 1236 | 80 |
| **BT** | A | 0.18 | 0.04 | | | | | | | 9.74 | 0.46 |
| **FY** | A | 0.19 | 0.04 | | | | | | | 9.23 | 0.44 |

[1] Vitezica ZG, Legarra A, Toro MA, Varona L. Orthogonal Estimates of Variances for Additive, Dominance, and Epistatic Effects in Populations. Genetics [Internet]. 2017 Jul [cited 2018 Jan 29];206(3):1297–307. Available from: http://www.ncbi.nlm.nih.gov/pubmed/28522540

**Table S3.3:** Literature review for inbreeding depression in some species of aquaculture. The inbreeding depression is expressed as the percentage decrease in the trait value per 10% increase in the inbreeding coefficient.

| Species | Trait | Inbreeding depression |
|---|---|---|
| Atlantic salmon[2] | BW | -0.6 to -2.6% |
| Rainbow trout[3] | BW | -2.3% |
| Rainbow trout[4] | BWH | -1.6 to -5.0% |
| Coho salmon[5] | BWH | -1.5% to -1.7% |

BW- Body Weight, BWH- Body Weight at Harvest

[2] Rye M, Mao IL. Nonadditive genetic effects and inbreeding depression for body weight in Atlantic salmon (Salmo salar L.). Livest Prod Sci. Elsevier; 1998;57(1):15–22.

[3] Hu G, Wang C, Da Y. Genomic heritability estimation for the early life-history transition related to propensity to migrate in wild rainbow and steelhead trout populations. Ecol Evol [Internet]. 2014 Apr 20 [cited 2015 Oct 16];4(8):1381–8. Available from: http://doi.wiley.com/10.1002/ece3.1038

[4] Pante MJR, Gjerde B, McMillan I. Effect of inbreeding on body weight at harvest in rainbow trout, Oncorhynchus mykiss. Aquaculture [Internet]. Elsevier; 2001 Jan 15 [cited 2018 Sep 4];192(2–4):201–11. Available from: https://www.sciencedirect.com/science/article/pii/S0044848600004671

[5] Neira R, Díaz NF, Gall GAE, Gallardo JA, Lhorente JP, Manterola R. Genetic improvement in Coho salmon (Oncorhynchus kisutch). I: Selection response and inbreeding depression on harvest weight. Aquaculture [Internet]. Elsevier; 2006 Jun 30 [cited 2018 Sep 4];257(1–4):9–17. Available from: https://www.sciencedirect.com/science/article/pii/S0044848606001839
